# PTEN Subcellular Localization Dictates Function

**DOI:** 10.64898/2026.06.16.732743

**Authors:** Nicole M. Desmet, Cheyenne F. Griffin, Helena Seo, Annaliese OuYang, Penelope Tir, Mackenzi L. Prina, Wei Wang, Meijie Li, Bryan W. Luikart

## Abstract

Mutations in phosphatase and tensin homolog (PTEN) drive unregulated activation of the phosphatidylinositol-3-kinase (PI3K) pathway, resulting in neuronal hypertrophy, and are strongly associated with autism spectrum disorder (ASD). Several PTEN mutations alter subcellular localization, yet how localization governs PTEN function in developing neurons remains unclear. Although PTEN has been reported broadly distributed throughout neurons, here, live imaging of HaloTagged PTEN reveals dynamically regulated localization, suggesting spatial control of its signaling. We then used retroviral-mediated genetic manipulation to delete endogenous *Pten* in developing hippocampal neurons while simultaneously expressing PTEN fused to defined localization motifs, allowing us to directly test how subcellular targeting regulates neuronal morphology. Loss of *Pten* produces neurons characterized by enlarged somata, more elaborate dendritic arbors, and increased spine density, length, and head area. Nuclear-excluded PTEN fully rescued these phenotypes, whereas targeting PTEN to filopodia via fusion to the FBAR domain of srGAP3 or to the postsynaptic density via Homer1C corrected or corrected all morphological abnormalities in PTEN-deficient neurons and simplified dendritic arborization compared to wild-type. In contrast, nuclear-localized PTEN produced only partial rescue, normalizing soma size and spine head area but not dendritic complexity or spine density. These findings indicate that PTEN acts locally to restrain growth and structural connectivity, whereas regulation of spine head size can be mediated by PTEN both inside and outside the nucleus, potentially through transcriptional or splicing-dependent mechanisms. Together, our results identify subcellular localization as a critical determinant of PTEN function and reveal spatially distinct mechanisms through which PTEN sculpts neuronal development.

## INTRODUCTION

Phosphatase and Tensin Homolog (PTEN), a protein and lipid phosphatase, downregulates the phosphatidylinositol-3-kinase pathway, a key signaling cascade regulating cell growth, metabolism, and survival. In the absence of PTEN, the pathway becomes overactive, leading to pathogenic growth, proliferation, and cytoskeleton rearrangement ^1–5^. Canonically acting at the membrane, PTEN also localizes to cytoplasmic organelles, including the nucleus, mitochondria, and endoplasmic reticulum, and can be secreted from cells ^6–9^. Furthermore, the subcellular localization of PTEN may dictate differential cellular function ^10–16^.

Various factors influence PTEN localization. PTEN’s C-terminal includes a C2 domain involved in membrane trafficking ^17^, a C-tail which, when phosphorylated, sequesters the protein away from the membrane in its “closed” conformation^18^, and a PDZ domain, which binds other proteins, stabilizing “open” PTEN and aiding its membrane binding^18^. The N-terminal contains the protein tyrosine phosphatase motif, a PIP_2-_binding motif which mediates binding to the membrane, a non-canonical nuclear localization sequence (NLS), and a cytoplasmic localization sequence. Mutations to the NLS motif inhibit nuclear entry^19,20^, while mutations in the cytoplasmic localization sequence lead to nuclear aggregation^20^. Additionally, SUMO1 modification can regulate PTEN’s association with the membrane^21^.

Nuclear PTEN maintains chromosomal integrity and regulates DNA repair ^22,23^. PTEN in the growth cone aids axonal guidance, cone collapse, and neurite growth ^24^. PTEN has also been documented within dendritic spine heads, and in the plasma membrane away from the post-synaptic density (PSD)^25^. Additionally, evidence suggests that during NMDAR-dependent plasticity, PTEN translocates and anchors in the PSD ^26^. Less is known about PTEN in dendritic processes, where it is hypothesized to regulate filopodial protrusion, retraction and persistence^1,27^.

PTEN mutations have been associated with Autism Spectrum Disorders (ASD). Most PTEN mutations associated with ASD do not fully abrogate PTEN’s phosphatase activity, suggesting a loss of catalytic activity may not be the only driving factor of autism-like phenotypes ^28–30^. Further studies of ASD-associated mutations found that many affect the subcellular localization of PTEN ^31^. Some mutants had decreased nuclear PTEN, and others had increased nuclear PTEN ^32^. Hence, the role of PTEN subcellular localization and its relevance to disease models is important.

Here, we examine PTEN in neurons and investigate the role of its subcellular localization in development. Utilizing conditional *Pten* knockout (*Pten^flx/flx^*) mice and retroviral infection, endogenous PTEN is replaced by expression of a PTEN-fusion protein to visualize and/or force localization of PTEN to various subcellular compartments. Our results indicate that PTEN localization is not ubiquitous throughout the cell, showing dynamic localization in developing neurons. Forced localization of PTEN to the nucleus results in modest regulation of soma size but dendrite growth and spine formation closely resemble neurons lacking PTEN entirely, suggesting that extra-nuclear PTEN is necessary to control growth of distal processes. Conversely, nuclear-excluded PTEN resulting in neurons with wildtype morphology suggest that dendritic spine density and length are under exclusive control of PTEN outside the nucleus. Neurons with PTEN localized to the filopodia or the post-synaptic density have decreased dendritic branching suggesting association with synapses and filopodia restricts dynamic dendrite growth and branching.

## RESULTS

### PTEN Localization is Regulated in Developing Neurons

We first imaged PTEN during the development of a granule neuron in the dentate gyrus of the hippocampus. We generated a retrovirus to simultaneously knockout endogenous *Pten* and express Halo9-tagged PTEN (pRubi-PTEN-Halo9-T2A-Cre). Retroviruses were injected into P7 *Pten*^flx/flx^xAi14 tdTomato Cre-reporter mice to express tdTomato, knockout endogenous *Pten* (as in ^33^), and express the PTEN-Halo9 fusion. We demonstrate that the PTEN-Halo9 is functional i*n vivo* by quantifying soma cross-sectional area of PTEN-Halo9 granule neurons and found they were not different than wildtype control neurons and significantly reduced in size compared to *Pten* KO neurons (**Fig. 1A-E**). We found that wildtype neurons infected with retrovirus to express GFP had typical soma cross-sectional area of 111.4±0.7716µm^2^ (**Fig. 1A,E**). When endogenous *Pten* was knocked out through Cre expression using the virus pRubiC-T2A-Cre, soma cross-sectional area more than doubles to 222.0±2.870µm^2^ (p<0.0001, **Fig. 1C,E**). When endogenous *Pten* is removed and replaced with human PTEN via the virus pRubi-PTEN-T2A-Cre, the neurons are smaller than wildtype neurons, likely due to the PTEN overexpression (96.34±1.662µm^2^, p<0.05, **Fig. 1B,E**). We next overexpressed GFP-T2A-PTEN ^31^ in wildtype neurons and found a trend towards decreased soma size that did not reach significance, likely due to a smaller sample size (78.79±3.887µm^2^, n=17, p=0.0647 vs wildtype, p=.9944 vs PTEN-T2A-Cre). When expressing PTEN-Halo9-T2A-Cre, soma size is not significantly different from wildtype (91.42±1.439µm^2^, p=0.0580). We performed live 2-photon imaging of *PTEN*^flx/flx^xAi14 mice infected with pRubi-PTEN-Halo9-T2A-Cre and were able to track PTEN’s localization in relation to the neuron’s growth. All PTEN-Halo9-positive neurons are positive for tdTomato, but we were unable to detect PTEN-Halo9 in some tdTomato positive neurons. The relative expression of tdTomato and PTEN-Halo9 varies so that some neurons may appear to be more red or cyan. Our results show that PTEN is not ubiquitously expressed and localized independently from tdTomato in regions throughout the cell, including dendritic growth cones and filopodia (**Fig. 1F-I, Supplemental Videos**). This demonstrates that PTEN localization is dynamically regulated.

**Figure 1.**
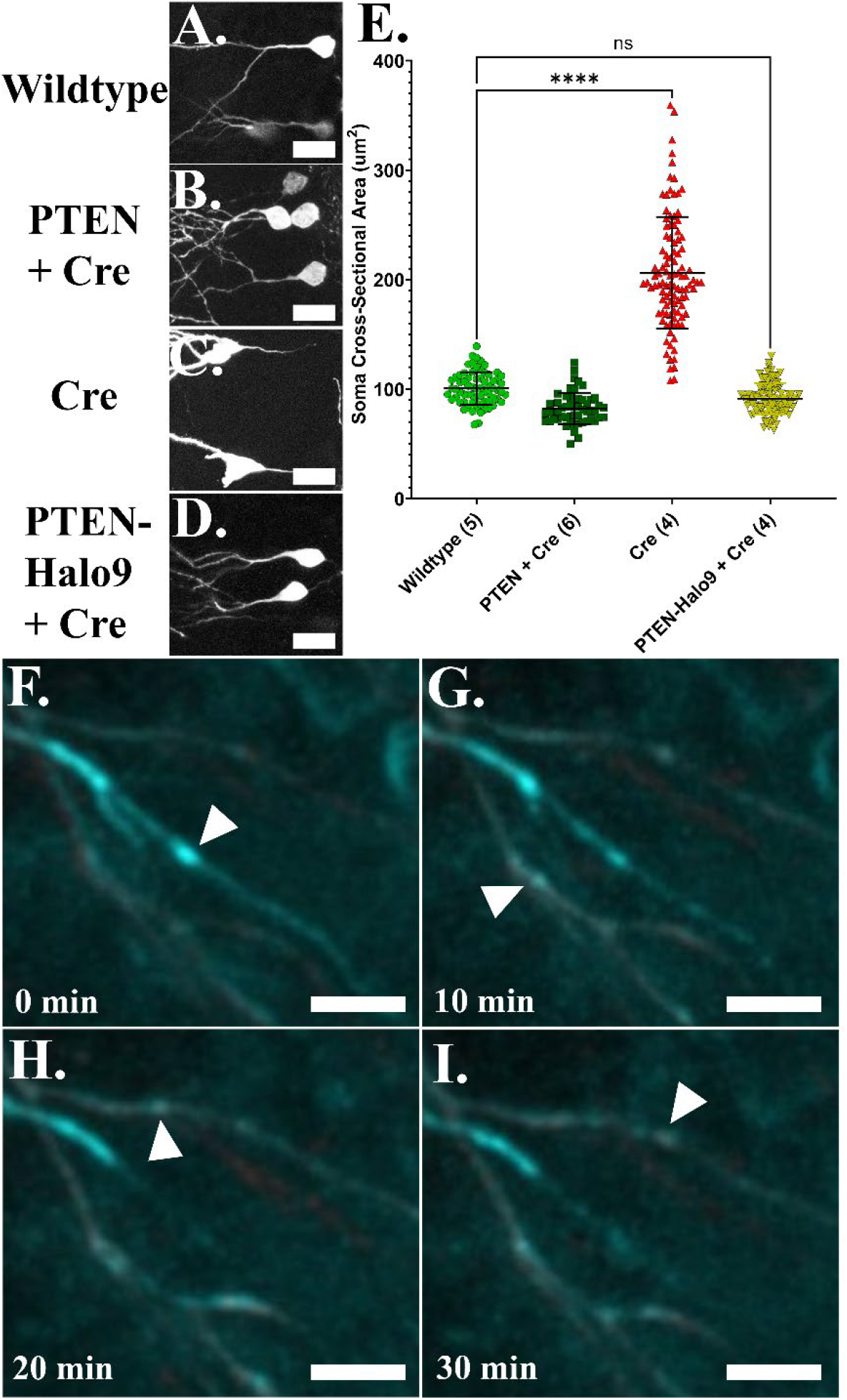
PTEN-Halo9 is functional in vivo and displays dynamic localization in growing neurons. P7 *Pten*^flx/flx^ x Ai14 mice are infected with pRubi (Wildtype; A.), PTEN-T2A-Cre (PTEN OE; B.), GFP-T2A-Cre (PTEN KO; C.), or PTEN-Halo9-T2A-Cre (PTEN-Halo9; D.) via retroviral stereotaxic injection. 21 days post-injection (DPI) morphology is assessed using immunohistochemical staining for GFP (wildtype) or tdTomato (Cre-positive). E. Soma cross-sectional area demonstrates that PTEN-Halo9 neurons display soma sizes indistinguishable from wildtype or neurons expressing untagged PTEN (ns, one-way ANOVA with Tukey’s post hoc comparison). F-G. P7 *Pten*^flx/flx^ x Ai14 mice are infected with PTEN-Halo9-T2A-Cre and acute slices are generated for live 2-photon imaging at 7-12 DPI. PTEN-Halo9 was visualized using JF669 during a time series of a recurring z-stack, and puncta are indicated with white arrows. A.-D. scale bar = 20µm. F.-I. scale bar = 10µm. n in parentheses indicates animal number.

### PTEN Lifetime Assessed Via HaloTag Pulse-Chase

Using DELTA, a fluorophore-based pulse-chase technique to measure protein lifetime *in vivo* ^34^, we calculated the lifetime of PTEN in granule neurons of the DG in mice. We injected AAV-DJ-hSyn.PTEN.Halo9.T2A.Cre into mice at post-natal day 7 (P7), and 21 days post-injection (DPI) we administered a “pulse” of HaloTag dye JFX673. Followed by a “chase” of JF552 at 6 different time points (1.5h, 8h, 24h, 48h, 72h, 96h). At the 1.5h chase we saw very little JF552 indicating that the pulse dose saturates the Halo9 sites. Consistent with previous experiments, we found a consistent value for PTEN lifetime but the 24h, 48h, 72h, and 96h (**Fig. 2A-F”**) timepoints reported consistent estimates (**Fig. 2G-H**). The average PTEN lifetime calculated from 2 mice at each of the 24h, 48h, 72h, and 96h chases was 2.50±0.14 days.

**Figure 2.**
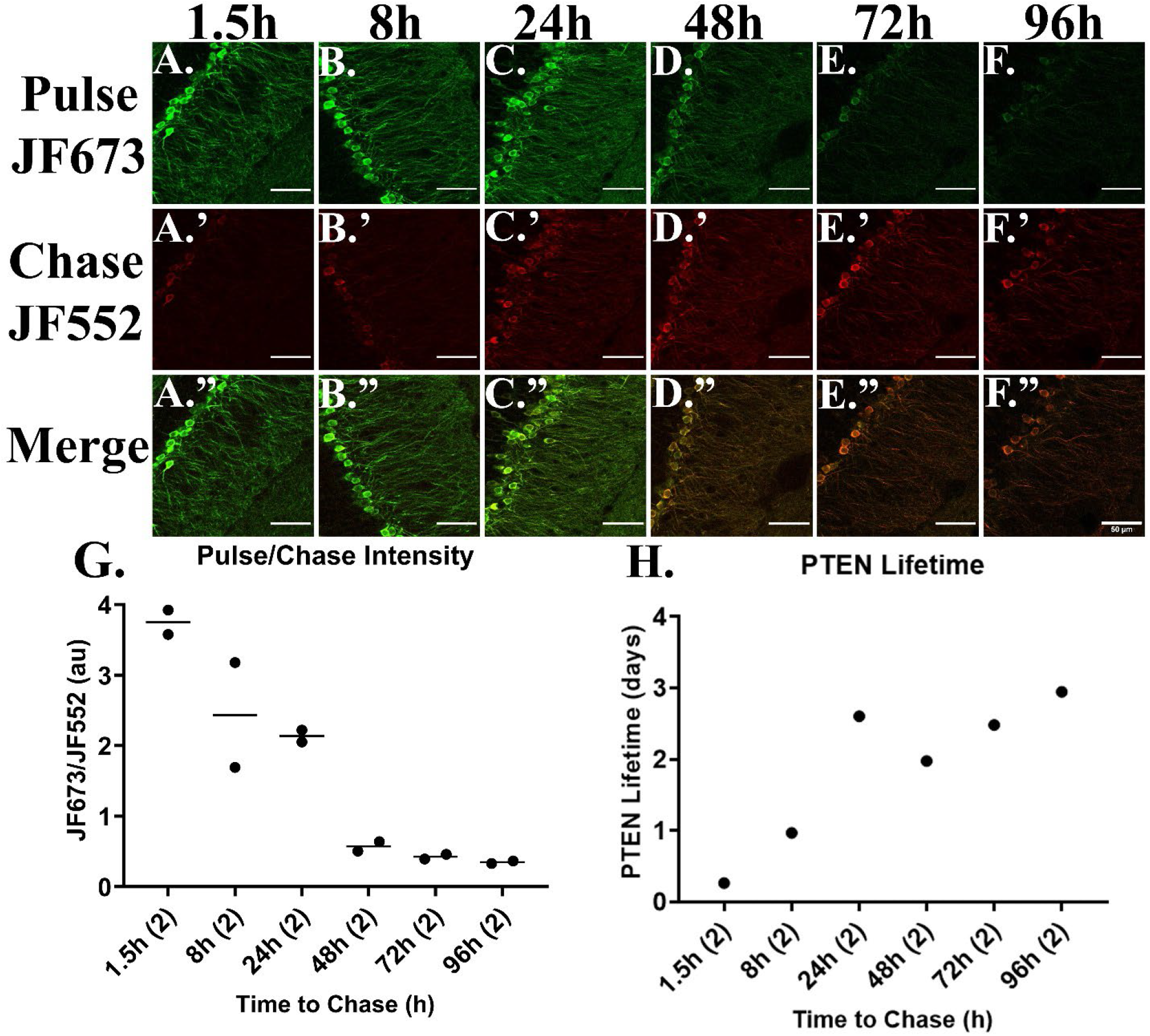
PTEN lifetime *in vivo*. **(A.-F.”)** Example multi-photon images from the granule cell layer and molecular layer in murine dentate gyrus, showing pulse, chase, and merged channels across all inter-pulse-chase time intervals assessed. **(G.)** Intensity of pulse (JF673) over intensity of chase (JF552) plotted against the pulse-chase time interval. Each point represents pulse/chase intensity of granule neurons in an individual mouse with lines representing the average for the time point. Pulse/chase intensity decreased as the time interval to the chase increased. **(H.)** PTEN lifetime plotted against the pulse-chase time interval. Each point represents the average PTEN lifetime across two mice assessed at each time point. The average PTEN lifetime calculated from the 24h, 48h, 72h, and 96h chases was 2.50±0.14 days.

### PTEN Localization is Controlled Using Retroviral-Mediated Genetic Manipulation

To determine the function of PTEN in different subcellular compartments we developed a series of retroviruses to knockout endogenous *Pten^flx/flx^* using Cre recombinase with PTEN fused to different domains to bias its localization to different subcellular compartments (**Fig. 3A**). To localize PTEN to the nucleus, two SV40 nuclear localization sequences (NLS) were fused to PTEN. To exclude PTEN from the nucleus, the Rev nuclear exclusion sequence (NES) was fused with PTEN. To localize PTEN to the filopodia, the FBAR domain of srGAP3 ^35^, was fused to PTEN. To localize PTEN to the post-synaptic density, PTEN was fused with HOMER1C, an adaptor protein found in the post-synaptic density. We used AlphaFold to predict the structures of these fusion proteins to display the relative size of the domains and display the predicted orientation of those domains to one another (**Fig. 3B**). Additionally, separate viruses were made with Halo9 fused to each of the PTEN fusion proteins.

**Figure 3.**
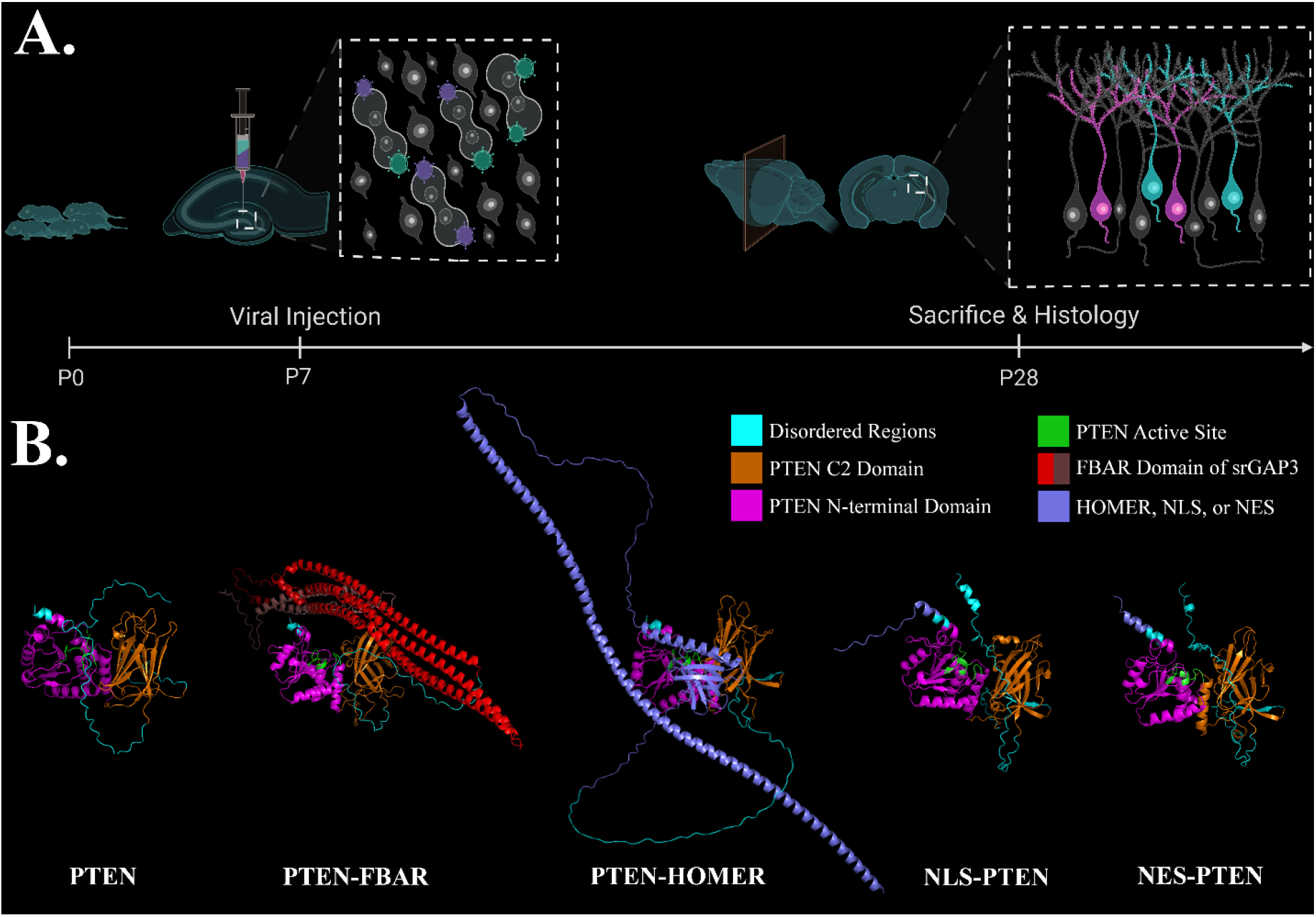
Experimental timeline and AlphaFold structure predictions for PTEN-fusion proteins. A. The dentate gyrus of P7 *Pten*^flx/flx^ x Ai14 (C57BL/6J) mice is stereotaxically injected with retroviruses. Viruses can include pRubi-GFP to label wildtype neurons, Cre to knockout endogenous *Pten* and elicit tdTomato reporter transgene, PTEN-T2A-Cre to reconstitute human PTEN, or PTEN fused to FBAR, HOMER, NLS or NES to direct the localization of PTEN to filopodia, post-synaptic densities, the nucleus and nuclear export, respectively. 21 days past injection (DPI) the mice are perfused and immunohistochemistry is used to examine neuronal morphology. B. AlphaFold predictions of the fusion proteins used to localize PTEN.

To assay whether these fusion proteins altered the subcellular localization of PTEN, we first visualized live HEK293ft cells infected with GFP-expressing and the various PTEN-Halo9 viruses stained with JF552 **(Fig. 4A-E.’)**. In HEK cells, NLS-PTEN-Halo9 and NES-PTEN-Halo9 displayed discrete nuclear localized and excluded expression. Qualitatively, the PTEN-FBAR-Halo9 appeared more filopodial while PTEN-HOMER was indistinguishable from wild type.

**Figure 4.**
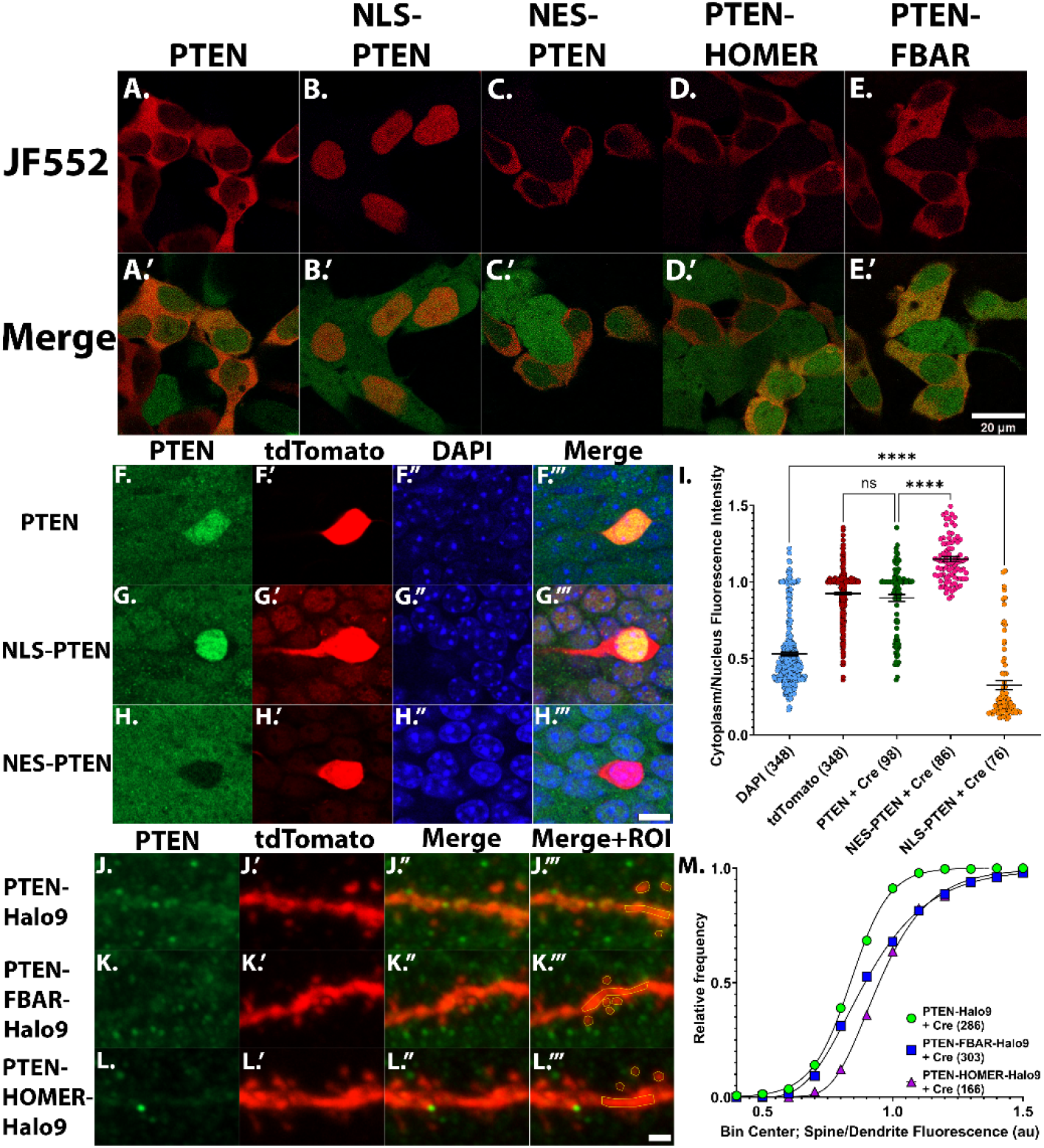
Fusion proteins alter PTEN subcellular localization. A-E. HEK293ft cells were infected with retroviruses expressing GFP (green) and PTEN-Halo9 fusion proteins were labeled with HaloTag ligand JF-552 (red). Confocal images of somas of dentate gyrus granule neurons of *Pten*^flx/flx^ x Ai14 mice infected with PTEN-T2A-Cre (F.-F.’’’), NLS-PTEN-T2A-Cre (G.-G.’’’), NES-PTEN-T2A-Cre (H.-H.’’’’), PTEN-Halo9-T2A-Cre (J.-J.’’’), PTEN-FBAR-Halo9-T2A-Cre (K.-K.’’’), or PTEN-HOMER-Halo9-T2A-Cre (L.-L.’’’). PTEN localization was quantified in regions of interest using the ImageJ polygon tool to compare fluorescence intensities of cytoplasm/nucleus or spine/dendrite. Cytoplasm/nucleus fluorescence intensity (I.) and cumulative probability distribution of spine/dendrite (M.) displayed at right. A.-E.’ scale bar = 20 µm. F.-H.’ scale bar = 20 µm. J.-L.’ scale bar = 5 µm. (Kolmogorov Smirnov test comparing frequency distribution of NLS-PTEN, NES-PTEN, PTEN-FBAR or PTEN-HOMER to PTEN **** = p < 0.0001), n in parentheses indicated number of cells (I.) or spines (M.) included.

To determine whether these fusion proteins altered the subcellular localization in neurons, we performed confocal microscopy on fixed, stained brain slices taken from a P28 mouse injected with our retroviruses at P7. Immunohistochemistry for tdTomato allowed us to visualize infected neurons, DAPI for nucleus, and PTEN immunohistochemistry allowed us to compare endogenous PTEN in the surrounding wildtype neurons to the exogenously expressed PTEN in tdTomato positive neurons. Previously, we quantified PTEN expression using both IHC and quantitative western blots after lentiviral expression (to infect all cells) using same ubiquitin promoter and all other elements within the ITRs as in the current retroviruses, showed a modest 1.75 fold overexpression of PTEN ^31^. Because our goal was to isolate the variable of subcellular localization with as close to endogenous expression levels as possible we performed all experiments with endogenous *Pten* deleted. We examined the fluorescence intensity of neurons expressing pRubi-PTEN-T2A-Cre compared to surrounding uninfected neurons. In the granule cell layer, PTEN fluorescence intensity of infected neurons is 77.48±3.592au, while uninfected neurons are 65.06±2.396au (p<0.01, **Fig. 4A-A’’’**). Qualitatively, NLS-PTEN neurons displayed prominent nuclear staining with staining absent in the surrounding cytoplasm (**Fig. 4B-B’’’**) and NES-PTEN had a lack of PTEN in the nucleus (**Fig. 4C-C’’’**). Quantitation of the cytoplasm/nucleus ratio of PTEN IHC fluorescence intensity compared to PTEN-Cre, DAPI, and tdTomato displayed bias of PTEN staining into the intended compartment. For PTEN the cytoplasm/nucleus fluorescence intensity was 0.8955±0.0225au, for NLS-PTEN the ratio was 0.3251±0.0298au (p<0.0001 vs PTEN, Table 1), and NES-PTEN the cytoplasm/nucleus ratio was 1.149±0.0172au (p<0.0001 vs PTEN, Table 1). For comparison, the cytoplasm/nucleus ratio for DAPI and tdTomato controls was 0.5297±0.0122au and 0.9248±0.0086au, respectively (**Fig. 4D**, Table 1).

**Table 1:** Statistical description of data presented in Figures.

| Condition | Mean±SEM<br>($\mu\text{m}^2$) | p-value<br>(vs WT) | n; cells<br>(animals) |
| --- | --- | --- | --- |
| Wildtype | 111.4±0.7716 | - | 447 (12) |
| PTEN + Cre | 96.34±1.662 | 0.0460 | 146 (6) |
| Cre | 222.0±2.870 | <0.0001 | 367 (5) |
| PTEN-Halo9 + Cre | 91.42±1.439 | 0.0580 | 99 (4) |

**Cytoplasm/Nucleus Fluorescence Intensity – Figure 4, graph not shown**
| Condition | Mean±SEM<br>(au) | p-value<br>(vs PTEN + Cre) | n; cells<br>(animals) |
| --- | --- | --- | --- |
| DAPI | 0.5297±0.0122 | <0.0001 | 347 (3) |
| tdTomato | 0.9248±0.0086 | 0.7143 | 347 (3) |
| PTEN + Cre | 0.8955±0.0225 | - | 85 (1) |
| NES-PTEN + Cre | 1.149±0.0172 | <0.0001 | 75 (1) |
| NLS-PTEN + Cre | 0.3251±0.0298 | <0.0001 | 97 (1) |

**Spine/Dendrite Fluorescence Intensity – Figure 4, graph not shown**
| Condition | Mean±SEM<br>(au) | p-value<br>(vs PTEN-Halo9) | n; spines<br>(cells) |
| --- | --- | --- | --- |
| tdTomato | 0.347±0.0071 | <0.0001 | 752 (46) |
| PTEN-Halo9 + Cre | 0.883±0.0077 | - | 285 (15) |
| PTEN-FBAR-Halo9 + Cre | 0.985±0.0131 | <0.0001 | 302 (15) |
| PTEN-HOMER-Halo9 + Cre | 1.020±0.0138 | <0.0001 | 165 (16) |

**Soma Area – Figure 4P.**
| Condition | Mean±SEM<br>($\mu\text{m}^2$) | p-value<br>(vs WT) | n; cells<br>(animals) |
| --- | --- | --- | --- |
| Wildtype | 107.6±0.9598 | - | 355 (12) |
| Cre | 222.0±2.870 | <0.0001 | 367 (5) |
| PTEN + Cre | 96.34±1.662 | 0.2140 | 146 (5) |
| PTEN-FBAR + Cre | 103.8±1.827 | 0.0070 | 125 (6) |
| PTEN-HOMER + Cre | 103.7±1.618 | <0.0001 | 148 (5) |
| NES-PTEN + Cre | 98.32±2.100 | 0.0500 | 116 (5) |
| NLS-PTEN + Cre | 135.8±2.413 | <0.0001 | 115 (6) |

**No. Primary Dendrites – Figure 5H.**
| Condition | Mean±SEM (#) | p-value<br>(vs WT) | n; cells<br>(animals) |
| --- | --- | --- | --- |
| Wildtype | 1.000±0.0000 | - | 20 (7) |
| Cre | 2.429±0.1402 | <0.0001 | 25 (6) |
| PTEN + Cre | 1.000±0.0000 | >0.9999 | 28 (6) |
| PTEN-FBAR + Cre | 1.000±0.0000 | >0.9999 | 22 (9) |
| PTEN-HOMER + Cre | 1.000±0.0000 | >0.9999 | 17 (4) |
| NES-PTEN + Cre | 1.000±0.0000 | >0.9999 | 19 (4) |
| NLS-PTEN + Cre | 2.346±0.2071 | <0.0001 | 26 (7) |

**Total Dendritic Length – Figure 5K.**
| Condition | Mean±SEM<br>($\mu\text{m}$) | p-value<br>(vs WT) | n; cells<br>(animals) |
| --- | --- | --- | --- |
| Wildtype | 1216±47.85 | - | 20 (7) |
| Cre | 1573±47.94 | <0.0001 | 25 (6) |
| PTEN + Cre | 1218±39.73 | >0.9999 | 28 (6) |
| PTEN-FBAR + Cre | 1079±43.92 | 0.9760 | 22 (9) |
| PTEN-HOMER + Cre | 1082±47.44 | 0.9980 | 17 (4) |
| NES-PTEN + Cre | 1276±55.33 | 0.9990 | 19 (4) |
| NLS-PTEN + Cre | 1435±63.83 | 0.5460 | 26 (7) |

**Spine Density – Figure 6H.**
| Condition | Mean±SEM<br>(spines/μm) | p-value<br>(vs WT) | n; cells<br>(animals) |
| --- | --- | --- | --- |
| Wildtype | 1.940±0.0681 | - | 130 (11) |
| Cre | 3.102±0.1198 | 0.0100 | 99 (4) |
| PTEN + Cre | 1.430±0.0866 | 0.9540 | 71 (4) |
| PTEN-FBAR + Cre | 1.506±0.0920 | 0.3770 | 30 (5) |
| PTEN-Homer + Cre | 1.232±0.0655 | 0.9120 | 60 (5) |
| NES-PTEN + Cre | 1.225±0.0723 | 0.1730 | 56 (4) |
| NLS-PTEN + Cre | 2.558±0.1347 | 0.2020 | 59 (4) |

**Spine Length – Figure 6I.**
| Condition | Mean±SEM<br>(μm) | p-value<br>(vs WT) | n; cells<br>(animals) |
| --- | --- | --- | --- |
| Wildtype | 0.9386±0.0107 | - | 130 (11) |
| Cre | 1.1790±0.0150 | <0.0001 | 99 (4) |
| PTEN + Cre | 0.8081±0.0149 | 0.2330 | 71 (4) |
| PTEN-FBAR + Cre | 0.9405±0.0250 | >0.9999 | 30 (5) |
| PTEN-HOMER + Cre | 0.9020±0.0184 | >0.9999 | 60 (5) |
| NES-PTEN + Cre | 0.9159±0.0176 | 0.9950 | 56 (4) |
| NLS-PTEN + Cre | 1.1180±0.0197 | <0.0001 | 59 (4) |

**Spine Area – Figure 6J.**
| Condition | Mean±SEM<br>(μm<sup>2</sup>) | p-value<br>(vs WT) | n; cells<br>(animals) |
| --- | --- | --- | --- |
| Wildtype | 0.2840±0.0043 | - | 130 (11) |
| Cre | 0.3356±0.0078 | 0.0310 | 99 (4) |
| PTEN + Cre | 0.2987±0.0082 | 0.9990 | 71 (4) |
| PTEN-FBAR + Cre | 0.2715±0.0045 | 0.8250 | 30 (5) |
| PTEN-HOMER + Cre | 0.3058±0.0081 | 0.8380 | 60 (5) |
| NES-PTEN + Cre | 0.2900±0.0065 | 0.9980 | 56 (4) |
| NLS-PTEN + Cre | 0.2726±0.0055 | 0.9860 | 59 (4) |

**Spine Area – Figure 6N.**
| Condition | Mean±SEM<br>(μm<sup>2</sup>) | p-value<br>(vs WT) | n; cells<br>(animals) |
| --- | --- | --- | --- |
| Wildtype | 0.4456±0.1882 | - | 391 (5) |
| Cre | 0.6765±0.3272 | <0.0001 | 377 (3) |
| PTEN + Cre | 0.4776±0.2274 | 0.7940 | 182 (4) |
| PTEN-FBAR + Cre | 0.4598±0.2080 | 0.8210 | 209 (5) |
| PTEN-HOMER + Cre | 0.4715±0.2218 | 0.7210 | 194 (4) |
| NES-PTEN + Cre | 0.5421±0.2253 | 0.9530 | 309 (4) |
| NLS-PTEN + Cre | 0.4299±0.2284 | >0.9999 | 442 (6) |

**Spine Area – Figure 6O.**
| Condition | Mean±SEM<br>(μm<sup>2</sup>) | p-value<br>(vs WT) | n; cells<br>(animals) |
| --- | --- | --- | --- |
| Wildtype | 0.3043±0.1235 | - | 539 (5) |
| Cre | 0.3811±0.1768 | 0.0120 | 477 (3) |
| PTEN + Cre | 0.2997±0.1027 | 0.9030 | 355 (4) |
| PTEN-FBAR + Cre | 0.3195±0.1099 | 0.0960 | 347 (4) |
| PTEN-HOMER + Cre | 0.2747±0.1095 | 0.0420 | 396 (2) |
| NES-PTEN + Cre | 0.3081±0.1088 | 0.5740 | 633 (3) |
| NLS-PTEN + Cre | 0.3159±0.1217 | 0.0760 | 516 (5) |

In the molecular layer, it was difficult to distinguish PTEN overexpression from infected dendrites compared to PTEN from surrounding wildtype cells (**Fig 4F-F’’’**). Quantitatively, PTEN fluorescence intensity of infected dendrites was 61.59±4.566au, not statistically different from the PTEN fluorescence intensity of an ROI in the molecular layer adjacent to the dendrite, 52.20±4.138au (p=0.1398). Qualitatively, a change in PTEN localization was observed in PTEN-FBAR-Halo9 (**Fig. 4G-G’’’**) and PTEN-HOMER-Halo9 neurons (**Fig. 4H-H’’’**). We created a region of interest in dendritic spines and the dendritic shaft using the tdTomato channel and quantified the fluorescence intensity ratio of either tdTomato or PTEN IHC, demonstrating a shift in the cumulative probability of expression into the spines for both PTEN-HOMER-Halo9 and PTEN-FBAR-Halo9 neurons when compared to PTEN-Halo9 neurons. For PTEN-Halo9, the spine/dendrite fluorescence intensity ratio of PTEN was 0.883±0.0077au. For PTEN-FBAR-Halo9 neurons, the spine/dendrite fluorescence intensity ratio was 0.985±0.0131au, and for PTEN-HOMER-Halo9 the ratio was 1.020±0.0138au. To display this shift in localization we generated a cumulative probability plot for spine/dendrite ratios of individual spines seen in **Fig. 4I**. Kolmogorov-Smirnov tests reveal a statistical difference between the cumulative probability plots of PTEN-Halo9 and PTEN-FBAR-Halo9 neurons (p<0.0001), as well as between PTEN-Halo9 and PTEN-HOMER-Halo9 neurons (p<0.0001). These data demonstrate that the HOMER and FBAR fusions result in a modest bias of PTEN localization into dendritic protrusions.

### Nuclear PTEN Partially Regulates Soma Hypertrophy

Next, we investigated whether PTEN subcellular localization regulated soma growth. *Pten* KO dentate granule neurons display soma cross-sectional areas approximately twice that of wildtype neurons *(Pten* KO=222.0±2.870µm^2^ vs wildtype=111.4±0.7716µm^2^, p<0.0001, **Fig. 5A,P**, **Table 1**). Overexpression of human PTEN in *Pten* KO cells (**Fig. 5B**, **Table 1**) results in soma cross-sectional area similar to wildtype neurons (96.34±1.662µm^2^, p=0.2140 **Fig. 5B,P**, **Table 1**). Both PTEN-FBAR (103.8±1.827µm^2^, p=0.0070 vs wildtype) and PTEN-HOMER (103.7±1.618um^2^, p<0.0001 vs wildtype) expression recue soma cross-sectional area (**Fig. 5D-F,P, Table 1**). NES-PTEN expression also results in rescue of soma cross-sectional area to wildtype levels (98.32±2.100µm^2^, p=0.0500 **Fig. 5J-L,P, Table 1**). However, NLS-PTEN results in an intermediate soma size (135.8±2.413µm^2^) larger than wildtype neurons (p<0.0001) but smaller than *Pten* KO neurons (p<0.0001, **Fig. 5M-O,P, Table 1**).

**Figure 5.**
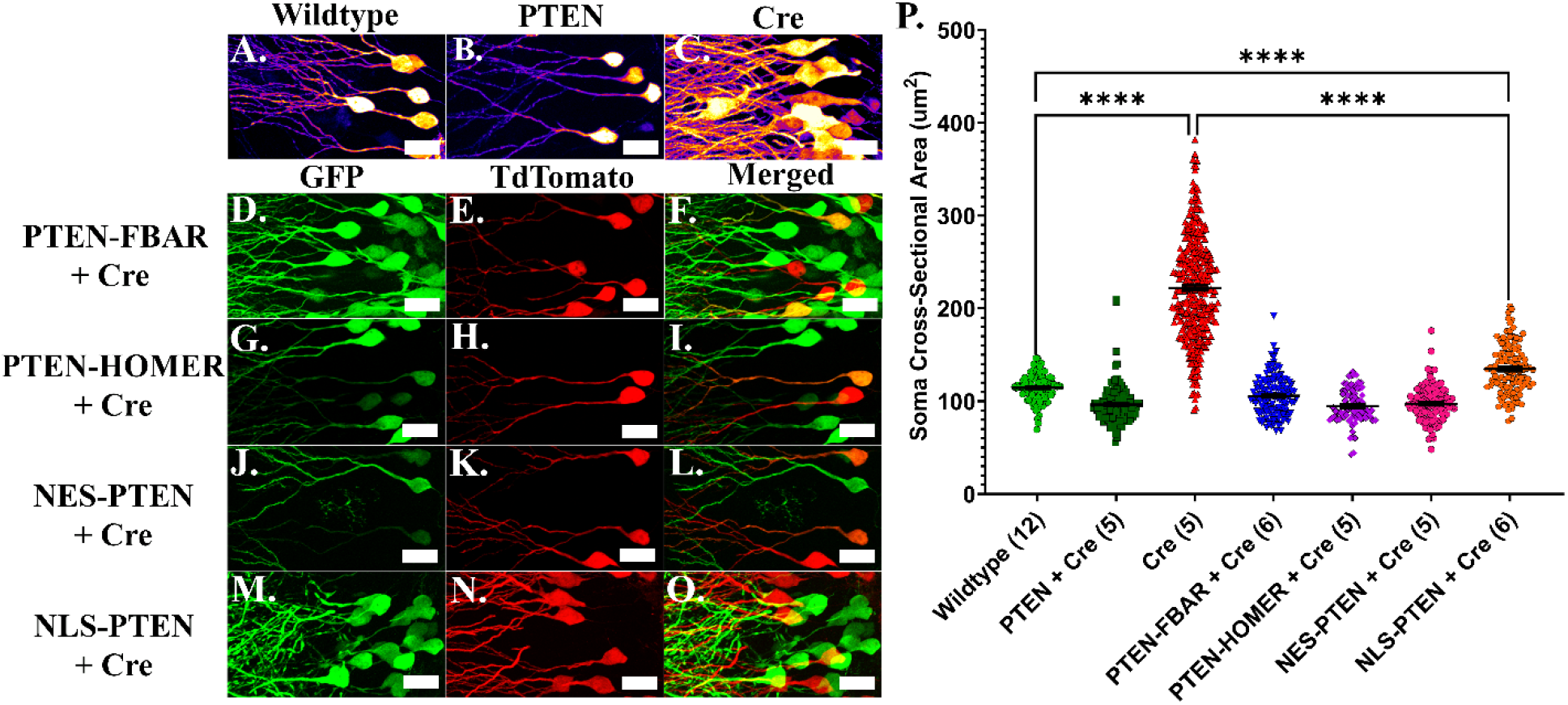
Nuclear PTEN Fails to Rescue Soma Hypertrophy of PTEN KO Neurons Confocal images of 21-day old granule neurons in the dentate gyrus of the hippocampus in PTEN^flx/flx^xAi14 mice. A-C. The ImageJ fire lookup table used to display 21DPI neurons from P7 mice were injected with pRubi-GFP (A. Wildtype), PTEN-T2A-Cre (B. PTEN reconstitution), or GFP-T2A-Cre (C. *Pten* KO). D-O. To examine the effect of altered subcellular localization PTEN^flx/flx^xAi14 mice were co-injected with pRubi-GFP (D, G, J, M. GFP, green in-tissue Wildtype control) and either PTEN-FBAR (E. tdTomato, red), PTEN-HOMER (H.), NES-PTEN (K.), or NLS-PTEN (N.). P. Soma cross sectional area was measured. PTEN, PTEN-FBAR, PTEN-Homer and NES-PTEN completely rescue somal hypertrophy seen in PTEN KO cells (Cre), while NLS-PTEN only partially rescues. Statistics performed using GraphPad Prism one-way ANOVA model with Tukey’s post hoc comparison, n in parentheses indicates the number of animals included. Scale bar = 20 µm. (* = p < 0.05, ** = p < 0.01, *** = p < 0.001, **** = p < 0.0001)

### Nuclear PTEN Does Not Regulate Dendritic Arborization

Next, we characterized the morphology of neurons with different PTEN subcellular localization patterns via sholl analysis. With Neurolucida 360, confocal images were used to reconstruct neurons in 3D (**Fig. 6A-G’**). Using retroviruses *in vivo*, we compare *Pten* KO neurons, wildtype neurons, and neurons in which endogenous *Pten* is knocked out and replaced with human PTEN bearing varying subcellular localization motifs. *PTEN* KO neurons have an increased number of primary dendrites (2.429±0.1402, p<0.0001 vs wildtype, Brown-Forsythe and Welch ANOVA with Dunnett’s, **Fig. 6A-H**, **Table 1**), altered dendritic architecture assayed by Sholl analysis (**Fig. 6I-L**), and increased total dendritic length (1573±47.94µm, p<0.0001, **Fig. 6K-M**, **Table 1**) when compared to wildtype neurons. We found that wildtype neurons and neurons expressing human PTEN had nearly identical arborization patterns throughout the granule cell layer, the inner molecular layer, the middle molecular layer, and the outer molecular layer (confirming ^3^) (**Fig. 6C**). There was no statistical difference between their primary dendrite averages (1.000±0.000 vs 1.000±0.000, p>0.9999, Brown-Forsythe and Welch ANOVA with Dunnett’s,), Sholl analyses, and total dendritic lengths (1218±40.85µm vs 1216±47.85µm, p>0.9999, **Fig. 6C,H-M, Table 1**). Neurons expressing NES-PTEN were also identical to wildtype neurons and those overexpressing exogenous human PTEN in their number of primary dendrites (1.000±0.0000, p>0.9999, Brown-Forsythe and Welch ANOVA with Dunnett’s), Sholl analysis, and total dendritic length (1276±55.33µm, p=0.8841, **Fig. 6F,H,J,M**). Neurons expressing NLS-PTEN appeared similar to PTEN KO neurons. NLS-PTEN neurons had an increase in number of primary dendrites (2.346±0.2071, p<0.0001, Brown-Forsythe and Welch ANOVA with Dunnett’s, **Fig. 6H**, **Table 1**), branching in the inner and middle-molecular layer in the Sholl analysis (**Fig. 6J**), and a trend towards an increase in total dendritic length (1435±63.83µm, **Fig. 6M**, **Table 1**) when compared to wildtype neurons. Individual means+/-SEM, p-values, and n numbers are summarized in Table 1. These results indicate that nuclear PTEN cannot fully regulate dendritic arborization.

**Figure 6.**
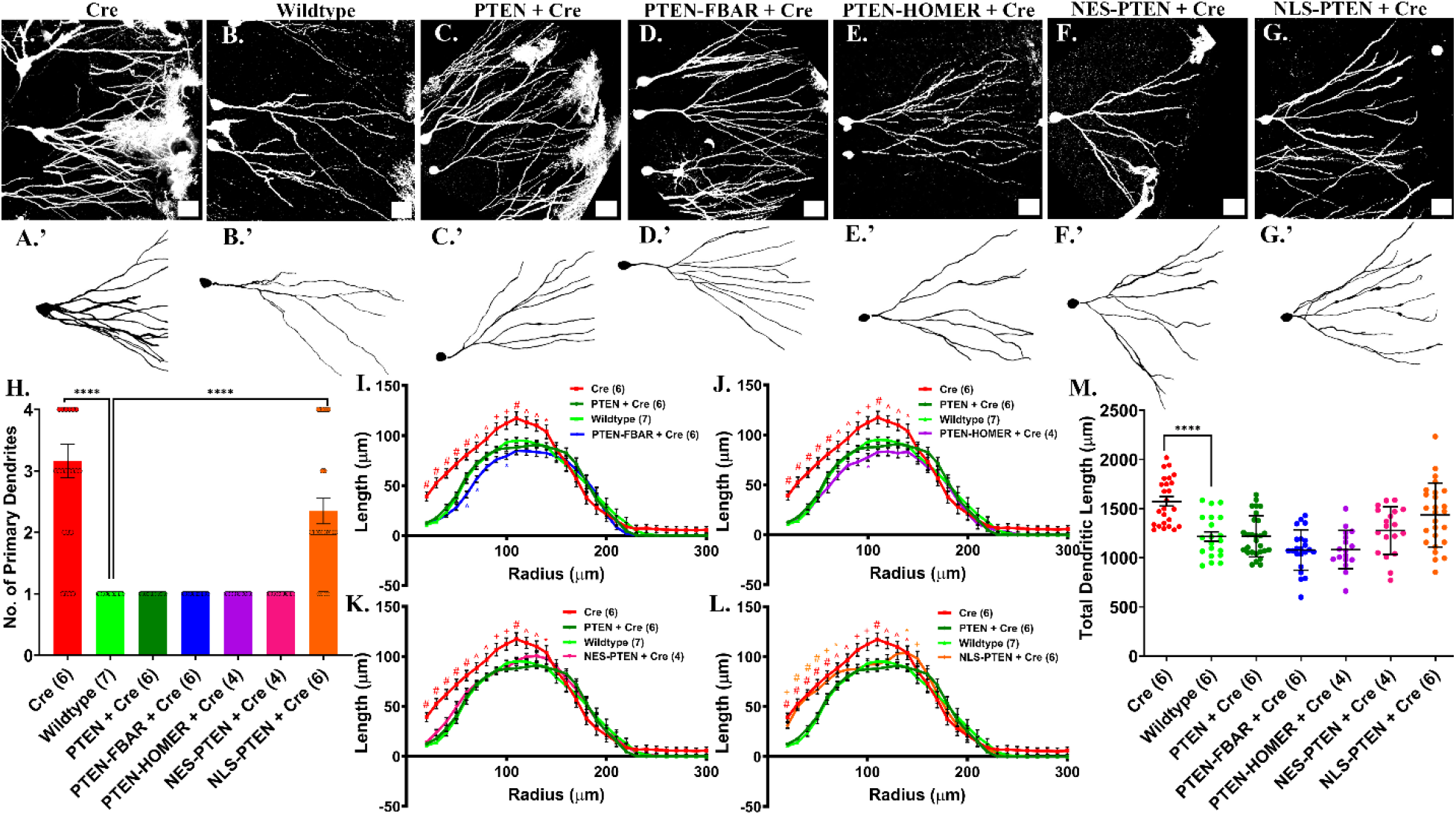
PTEN-FBAR and PTEN-HOMER decrease dendritic complexity and NLS-PTEN fails to rescue dendritic hypertrophy of PTEN KO neurons. A-G’. Confocal maximum projection images and NeuroLucida360 reconstructions of representative 21-day old neurons for each viral construct expressed in *Pten*^flx/flx^xAi14 mice. PTEN KO (Cre, A., A.’), Wildtype (B., B.’), human PTEN reconstitution (PTEN + Cre, C., C.’), PTEN-FBAR (D., D.’), PTEN-HOMER (E., E.’), NES-PTEN (F., F.’), and NLS-PTEN (G., G.’). H. Primary dendrite counts indicate that all constructs other that NLS-PTEN have a single primary dendrite arising from the soma. I-L. Sholl analysis displaying dendritic architecture for Wildtype (neon green) and *Pten* KO (Red, Cre) and human PTEN reconstitution (dark green) compared to PTEN-FBAR (blue, I.), PTEN-HOMER (Purple, J.), NES-PTEN (magenta, K.), and NLS-PTEN (Orange, L.). PTEN-FBAR and PTEN-HOMER have a decrease in branches in the regions between 50 and 100um from the soma while NLS-PTEN displays an increase in branches in the inner molecular layer comparable to that of PTEN KO neurons. NES-PTEN is equivalent to Wildtype. M. Total dendritic length also shows that NES-PTEN and PTEN result in wildtype arborization, PTEN-FBAR and PTEN-HOMER have decreased arborization and NLS-PTEN sows dendritic hypertrophy. All tracings are uploaded to neuoromorpho.org. H., M. statistics performed using GraphPad Prism Brown-Forsythe and Welch ANOVA with Dunnett’s multiple comparison test. I.-L. statistics performed using GraphPad Prism two-way ANOVA mixed-effects model. Scale bar = 25 µm. (I.-L. : * = p≤0.05, ^ = p≤0.01, + = p≤0.001, # = p≤0.0001) (H.,M. : * = p < 0.05, ** = p < 0.01, *** = p < 0.001, **** = p < 0.0001). n in parentheses indicates number of animals included.

### Filopodial and PSD-localized PTEN Decrease Dendritic Arborization

Neurons expressing PTEN-FBAR (**Fig. 6D**) or PTEN-HOMER (**Fig. 6E**) with PTEN localized to the filopodia or the PSD had a single primary dendrite (1.000±0.0000, p>0.9999, **Fig. 6H**, **Table 1**) and were less arborized in their Sholl analyses in the inner and middle molecular layer (p<0.0001 vs wildtype, **Fig. 6I,J**). The total dendritic length also showed a trend towards being lower than that of wildtype neurons and neurons expressing human PTEN. PTEN-FBAR neurons had a total dendritic length of 1079±43.92µm (p=0.9760 compared to wildtype, p<0.0001 compared to *Pten* KO). PTEN-HOMER neurons had a total dendritic length of 1082±47.44µm (p=0.9980 compared to wildtype, p<0.001 compared to *Pten* KO (Table 1). These results indicate that biasing PTEN localization towards the filopodial or synaptic compartments decreases dendritic arborization.

### Extra-Nuclear PTEN Rescues Dendritic Spine Density, Length, and Head Area While Nuclear PTEN Rescues Only Head Area

Lastly, we evaluated the effects of various PTEN subcellular localizations on dendritic spines in the middle molecular layer. For wildtype neurons, there are approximately 1.940±0.0681 spines/µm, which are 0.9386±0.0107µm long on average, and head cross-sectional area of these spines is 0.2840±0.0043µm^2^. In neurons lacking *Pten* (**Fig. 7C**), spine density (3.102±0.1198spines/µm, p=0.0100), length (1.1790±0.0150µm, p<0.0001), and head area (0.3356±0.0078µm^2^, p=0.0310) are increased (**Fig. 7H-J**, **Table 1**) compared to wildtype neurons (**Fig. 7A**). When PTEN is overexpressed in *Pten* KO cells (**Fig. 7B**), spine density (1.430±0.0866spines/µm, p=0.9540), length (0.8081±0.0149µm, p=0.2330), and spine head area (0.2987±0.0082µm^2^, p=0.9990) are equivalent to that of wildtype neurons (**Fig. 7H,I**, **Table 1**). The spine density results for PTEN-FBAR (1.506±0.0020spines/µm, p=0.9540, **Fig. 7D**), PTEN-HOMER (1.232±0.0655spines/µm, p=0.3770, **Fig. 7E**), and NES-PTEN (1.225±0.0723spines/µm, p=0.1730, **Fig. 7F**) were equivalent to what was seen for wildtype neurons (**Fig. 7H-J**, **Table 1**). The spine length results for PTEN-FBAR (0.9405±0.0250um, p>0.9999, **Fig. 7D**), PTEN-HOMER (0.9020±0.0184um, p>0.9999, **Fig. 7E**), and NES-PTEN (0.9159±0.0176um, p=0.9950 **Fig. 7F**) were also equivalent to what was seen for wildtype neurons (**Fig. 7H-J**, **Table 1**). Neurons expressing NLS-PTEN, however, display increased spine length (1.1180±0.0197µm, p<0.0001), and a trend towards increased spine density (2.558±0.1347spines/µm, p=0.2020) when compared to wildtype (**Fig. 7H,I**, **Table 1**). Interestingly, NLS-PTEN returned spine head area to wildtype values (0.2726±0.0055µm^2^, p=0.9860, **Fig. 7J**, **Table 1**). To confirm this result, we repeated the experiment with a separate set of animals using a different method. Spine heads were manually circled using 2-photon image stacks of 21-day old neurons from each group, images were thresholded, and spine head cross-sectional area was measured (**Fig. 7K-M**). Using this independent methodology on a second set of animals we reproduced the result demonstrating all PTEN localizations result in rescue to wildtype levels of spine head cross-sectional area, including NLS-PTEN (0.4299±0.2284um^2^ vs wildtype 0.4456±0.1182um^2^, p>0.9999, **Fig. 7N**, **Table 1**), while *Pten* KO neurons have an increased spine head cross-sectional area (0.6765±0.3272um^2^, p<0.0001). In addition to thresholding to determine the margins of the spine head, the perceived margin of spine heads were manually circled by a blinded investigator and cross-sectional area was measured (**Fig. 7O**). Again, *Pten* KO neurons have significantly increased spine head cross-sectional area when compared to wildtype neurons (0.3811±0.1768um^2^ vs 0.3043±0.1235um^2^, p=0.0120, **Fig. 7O**, **Table 1**). This increase was not seen for NLS-PTEN. In this analysis, PTEN-HOMER neurons had significantly smaller spine heads compared to wildtype neurons (0.2747±0.1095um^2^ vs wildtype, p=0.0420, **Fig 7O**, **Table 1**).

**Figure 7.**
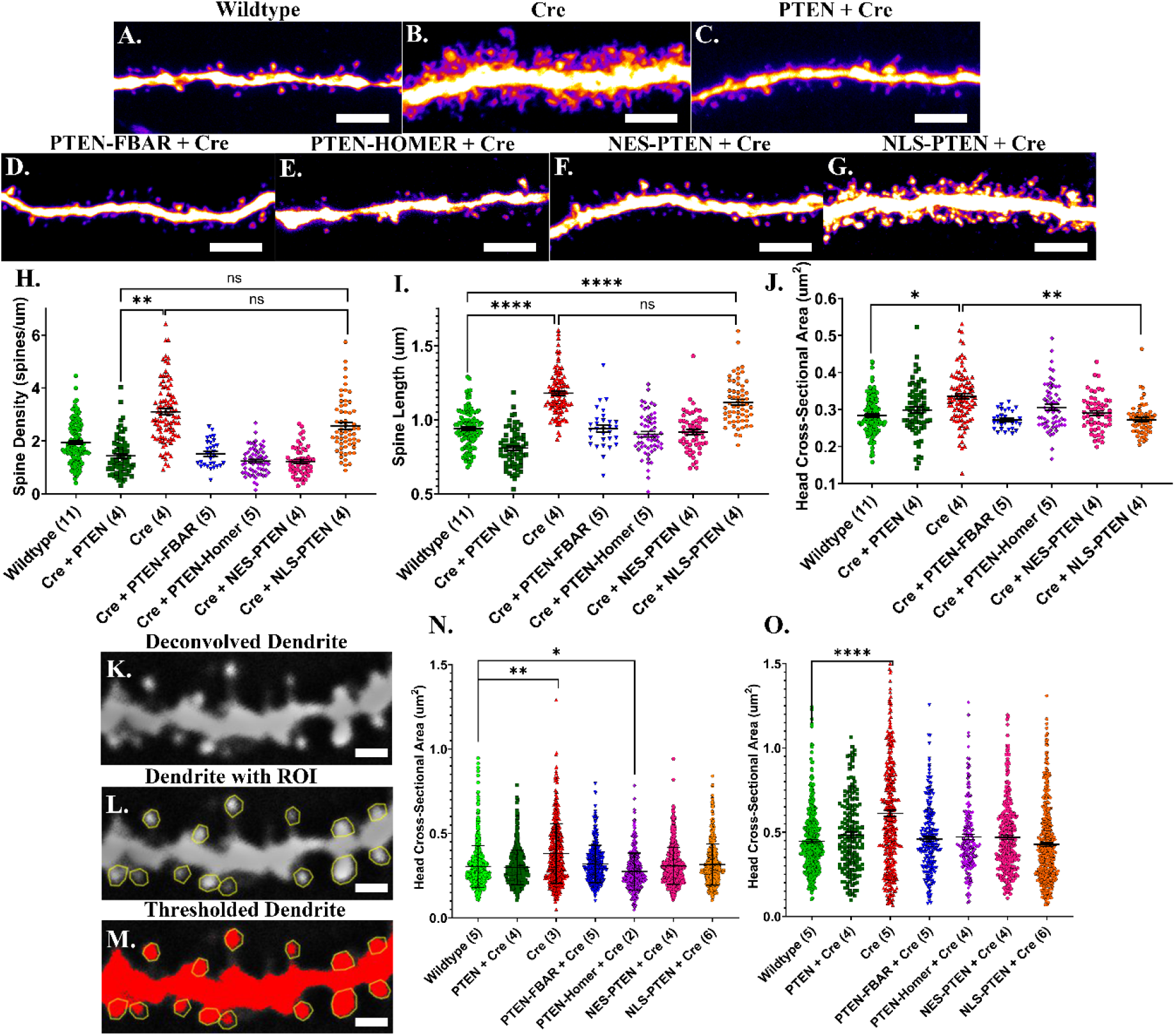
Extra-Nuclear PTEN rescues dendritic spine density, length, and head area while nuclear PTEN rescues only head area. A-G. Confocal maximum projection images with the ImageJ fire lookup table for 21-day old dentate granule neurons in PTEN^flx/flx^xAi14 mice infected with pRubi-GFP (A. Wildtype), PTEN KO (Cre, B.), human PTEN reconstitution (PTEN + Cre, C.), PTEN-FBAR (D.), PTEN-HOMER (E.), NES-PTEN (F.), and NLS-PTEN (G.). Quantitative results of dendritic spine density (H.), length (I.), and head area (J.) are displayed below. PTEN-FBAR, PTEN-Homer, and NES-PTEN completely rescue dendritic spine density, length, and head area, while NLS-PTEN only rescues the dendritic spine head area increases seen in PTEN KO cells. This result was confirmed using 2-photon z-stacks manually circling spine heads and thresholding to quantify spine head area based on standardized pixel intensity (K.-N.). We also manually circled the spine heads determining the area based on appearance rather than intensity threshold (O). Statistics performed using GraphPad Prism one-way ANOVA model with Tukey’s post hoc comparison. A.-G. scale bar = 5 µm. K.-M. scale bar = 2 µm (* = p < 0.05, ** = p < 0.01, *** = p < 0.001, **** = p < 0.0001). n in parentheses indicates number of animals included.

## DISCUSSION

PTEN is a critical regulator of neuronal growth and synaptic development, and retrovirus-mediated deletion of *Pten* in dentate gyrus granule neurons produces well-characterized phenotypes including soma hypertrophy, excessive dendritic elaboration, and increased dendritic spine density, length, and head area ^36–38^. These structural changes promote the formation of additional excitatory synapses and are predicted to alter both local and global integration of neuronal signals, potentially contributing to circuit dysfunction associated with neurodevelopmental disorders such as autism spectrum disorder ^39^. Despite this established role for PTEN in constraining neuronal growth, it has remained unclear how the subcellular localization of PTEN influences these processes. PTEN mutations in patients have been demonstrated to alter not only the phosphatase activity but also the localization of PTEN in both ASD ^31,32,40^ and cancer ^19,41,42^. Here we demonstrate that the impact of PTEN on neuronal development varies depending on its intracellular distribution.

### PTEN lifetime and localization

Using DELTA, a recently established Halo/fluorophore-based pulse-chase technique, we found that the average lifetime of PTEN in mouse hippocampal granule neurons was 2.50±0.14 days. The lifetime for GluA2, PSD-95, and MeCP2 were approximately 5.5, 14.5, and 12 days, respectively ^34^. Therefore PTEN lifetime is roughly half that of GluA2 and much shorter than PSD-95 and MeCP2. We interpret this to indicate that PTEN is relatively unstable and/or highly regulated. Live two-photon imaging revealed that PTEN dynamically moves in discrete compartments, including dendritic growth cones and filopodia. This observation is consistent with studies in motile cells showing that the relative distribution of PI3K and PTEN establishes localized phosphoinositide gradients that direct membrane expansion and cytoskeletal remodeling ^43,44^. Similar compartmentalized signaling has been proposed in neurons, where PIP3 enrichment promotes dendritic growth and synapse formation ^3,27,45^. We noted instances where the growth cone of one neuron collided with a dendrite shaft and the intensity of PTEN-Halo in the growth cone increased (see both supplemental videos). While this observation needs quantitation in the future, this could be the basis for the lack of dendrite self-avoidance after *Pten* knockout ^46^. Our findings demonstrate that PTEN displays punctate expression and is dynamically trafficked to growth-associated structures, placing it in position to spatially constrain PI3K-dependent signaling.

### Extra-nuclear PTEN is the primary regulator of soma growth

Deletion of *Pten* produced the expected soma hypertrophy, a phenotype widely attributed to hyperactivation of the PI3K–Akt–mTOR pathway. mTORC1 activation enhances cap-dependent translation, including that of 5′TOP mRNAs encoding ribosomal proteins and translational machinery, thereby driving cellular enlargement ^47–50^. The ability of nuclear-excluded PTEN to fully rescue soma size strongly suggests that suppression of this translational program occurs outside the nucleus. These data align with canonical models in which PTEN acts at the plasma membrane to dephosphorylate PIP3 and limit Akt activation upstream of mTOR.

The partial rescue observed with nuclear-restricted PTEN implies that transcriptional regulation by PTEN may contribute. Nuclear PTEN has been implicated in chromatin stability, DNA repair, and transcriptional control, processes that may influence neuronal function over longer timescales ^23,32,51^. It is also possible that some PTEN exits the nucleus although it is immediately trafficked back into the nucleus. If this is the case there may be a small degree of translation regulation still occurring in NLS-PTEN. Regardless, our results indicate that these nuclear roles are insufficient to counteract the potent growth-promoting effects of unchecked cytoplasmic PI3K signaling during development. Thus, regulation of protein synthesis rather than transcription appears to be the dominant mechanism controlling neuronal soma size.

### Nuclear PTEN is insufficient to regulate dendritic architecture

Dendritic arborization was similarly dependent on extranuclear PTEN. Whereas wild-type and cytoplasmic PTEN restored normal branching patterns, nuclear-restricted PTEN failed to prevent the increased primary dendrites and total dendritic length characteristic of *Pten*-deficient neurons. PI3K signaling is known to regulate cytoskeletal dynamics through downstream effectors including small GTPases and actin regulatory proteins, suggesting that access to cytoplasmic signaling networks is required for PTEN to shape dendritic structure ^25,52,53^. These findings support a model in which dendritic morphogenesis is governed by localized lipid signaling rather than nuclear transcriptional programs alone.

### Spatial enrichment of PTEN restricts dendritic elaboration

Targeting PTEN to filopodial, PTEN-FBAR, or postsynaptic PTEN-HOMER compartments modestly reduced dendritic complexity relative to wild-type neurons, suggesting that localized PTEN activity can suppress structural growth within discrete microdomains. This interpretation is consistent with studies of chemotaxis demonstrating that localized PTEN opposes PI3K-driven protrusion ^54,55^. Further, this supports a parallel regulation of these pathways in neurons to support filopodial and growth cone sprouting ^27,56,57^. We interpret the similarity of PTEN-FBAR and PTEN-HOMER in dendrite regulation to support the idea that filopodial outgrowth, synapse formation, and branch formation of growing dendrites and axons are interrelated ^58–62^. In neurons, such spatial regulation may allow branches to stabilize or retract in response to local signaling cues, providing a mechanism for fine control of dendritic architecture during circuit formation.

### Compartment-specific regulation of dendritic spines

PTEN loss increased spine density, length, and head size, consistent with enhanced excitatory synaptogenesis reported in models of PI3K pathway hyperactivation. Restoration of extranuclear PTEN normalized these parameters, whereas nuclear PTEN selectively rescued spine head size but not spine density or length. Optical measurements tend to overestimate the physical dimensions of submicron structures relative to EM reconstructions, which show dentate granule cell spine heads typically have diameters of approximately ∼0.45–0.55 µm, corresponding to cross-sectional areas near ∼0.15–0.30 µm² ^63,64^. Importantly, this is because the lateral resolution of conventional two-photon or confocal microscopy is ∼300–500 nm depending on numerical aperture and wavelength, and therefore optical measurements should be interpreted as relative differences between experimental groups rather than absolute spine size. Consequently, because spine head area correlates with postsynaptic strength, this result raises the possibility that nuclear PTEN contributes to aspects of synaptic maturation or neuronal activity, while dendritic PTEN limits synapse formation. This was a surprising finding and warrants future study of potential mechanisms like PTEN-dependent regulation of transcription ^65,66^ or alternative splicing ^67^. Such compartment-specific effects suggest that PTEN coordinates multiple stages of synaptic development through spatially distinct mechanisms.

### Implications for models of PTEN function in developing neurons

Several considerations qualify the interpretation of our localization experiments. Although fusion motifs effectively biased PTEN toward specific subcellular compartments, they do not confer absolute spatial restriction. PTEN remains a dynamically trafficked protein and may retain activity while in transit between compartments; thus, constructs such as PTEN-HOMER should not be interpreted as exclusively confined to dendritic spines. Even targeting strategies that appeared highly specific, including nuclear localization and export sequences, likely permit some degree of nucleocytoplasmic shuttling, indicating that our approach favors enrichment rather than precise spatial control. Future studies employing extended live imaging of HaloTagged PTEN will be important for defining the kinetics of PTEN trafficking and determining whether neuronal activity, growth factors, or other extracellular cues regulate its intracellular distribution. In addition, because the present work focuses primarily on structural outcomes, electrophysiological experiments will be necessary to establish how compartment-specific PTEN localization influences synaptic function and intrinsic excitability. Addressing these questions will further refine our understanding of how spatial regulation of PTEN contributes to neuronal development and circuit function.

Collectively, our findings support a model in which PTEN functions predominantly at extranuclear sites to antagonize PI3K–Akt–mTOR signaling and restrain neuronal growth. Although nuclear PTEN has been implicated in chromatin regulation and genome maintenance, our data indicate that its presence alone is not sufficient to counterbalance cytoplasmic growth signals during critical developmental windows. These observations suggest that the biological output of PTEN is governed not only by its enzymatic activity but also by its spatial distribution within the cell. This spatial framework has important implications for how neuronal morphology is established. Because pathways downstream of PTEN directly influence protein synthesis, cytoskeletal dynamics, and membrane expansion, even modest shifts in PTEN localization could bias the balance between growth and stabilization, thereby shaping dendritic architecture. We therefore speculate that the regulation of PTEN localization may represent a mechanism contributing to the morphological diversification of neurons during development.

## ACKNOWLEDGEMENTS

We would like to thank Thomas Sudhof and Xian Jiang for sharing the PTEN-HOMER1C fusion construct. We would like to thank Luke Lavis for generous sharing of JaneliaFlour-Halo dyes and Boaz Mohar for advice on their use *in vivo* and on acute slices. NMD would like to thank her thesis committee, Hermes Yeh, Robert Hill, and Mike Hoppa for valuable feedback.

## AUTHOR CONTRIBUTIONS

ML and WW generated tools for this study. NMD, HS, AO, MLP, PT, CFG, and BWL performed experiments. NMD and BWL performed analyses. NMD wrote the manuscript. BWL edited the manuscript. This work was funded by R01 (MH097949) to BWL. This work was also supported by Optical Cellular Imaging Shared Resource at the Geisel School of Medicine at Dartmouth College and the Civitian Cellular Imaging Core at the University of Alabama at Birmingham.

## CONFLICT OF INTEREST STATEMENT

All authors declare no biomedical financial interests or potential conflicts of interest.

## STAR METHODS

### KEY RESOURCES TABLE

| REAGENT or RESOURCE | SOURCE | IDENTIFIER |
| --- | --- | --- |
| Antibodies |  |  |
| Bacterial and virus strains |  |  |
| Experimental Models:<br>Organisms/strains |  |  |
| Deposited Data |  |  |
| Neuronal Reconstructions | Neurolucida360 | Neuromorpho.org: Luikart Archive |

### RESOURCE AVAILABILITY

#### LEAD CONTACT

For further information and any requests for resources and reagents, can be directed to and will be fulfilled by the lead contact, Bryan Luikart

#### MATERIALS AVAILABLILTY

All mice and viral constructs used in this study are all available through Addgene or Jackson laboratories with individual catalog numbers for each item listed in the key resources table and the subsections below. JFX dye was generously supplied by the Janilea Institute

#### DATA AND CODE AVAILABLILITY

NeuroLucida360 neuronal reconstruction data have been deposited at Neuromorpho.org and are publicly available as of the date of publication. Other data reported in this study consist of image stacks, ROI sets for image analysis, spreadsheets for quantitative image analysis.

To request access, contact the lead contact. This paper does not report original code. Any of this data or additional information required to reanalyze the data reported in this paper is available from the lead contact upon request.

### EXPERIMENTAL MODEL AND SUBJECT DETAILS

#### ANIMALS

All procedures were approved by the Dartmouth Institutional Animal Care and Use Committee (IACUC), the Association for Assessment and Accreditation of Laboratory Animal Care Review Board, and the University of Alabama at Birmingham IACUC. Mice of both sexes were used. *Pten^flox/flox^* (B6.129S4-*Pten^tm1Hwu^*/J) mice were backcrossed into the C57BL/6J background at least five generations. *Pten^Flox/Flox^* x tdTomato (Ai14)*^Flox/Flox^* mice were generated by crossing *Pten^Flox/Flox^* (B6.129S4-*Pten^tm1Hwu^*/J) mice to tdTomato (Ai14)*^Flox/Flox^* (B6;129S6-*Gt(ROSA)26Sor^tm14(CAG-TdTomato)Hze^*/J) mice. Animals were provided food and water *ad libitum* and housed on a 12-hour light/dark cycle.

### METHOD DETAILS

#### REPLICATION-DEFICIENT RETROVIRUSES

Retroviral packaging was performed as described in^68^.

Replication deficient retroviruses utilized the pRubi construct described in^69^. The pRubi-Pten-T2A-Cre was previously generated^3^. All viral constructs are being provisionally submitted to addgene. For the pRubi-PTEN-HALO-T2A-Cre, the Halo9 tag was cloned from pcDNA5/FRT/TO_H2B-HaloTag7-T2A-Tom20-HaloTag9 (addgene 175547)^70^ with the following Gibson reaction:

pRubi-Pten-2a-cre_fwd aatctccggtTCTAGAGAGGGCAGAGGAAG pRubi-Pten-2a-cre_rev cgctaccgccGACTTTTGTAATTTGTGTATGCTGATC Halo9_fwd tacaaaagtcGGCGGTAGCGGGGATCCA Halo9_rev cctctctagaACCGGAGATTTCCAGGGTTGAC

The quickchange lightning mutagenesis kit (Agilent 210519) was used to generate pRubi-NLS-PTEN-T2A-Cre and pRubi-NES-PTEN-T2A-Cre. We inserted 2x SV40 NLS sequences with the following mutagenesis oligo: ggtcgactctagaaccatgCCTAAGAAGAAGAGGAAGGTACCGGTCGCCACCATGCCTAAGAAGAA GAGGAAGGTGacagccatcatcaaagagatcgpRubi-NES-PTEN-T2A-Cre was generated by inserting the Rev NES with the following: gcttgggctgcaggtcgactctagaaccatgCTGCCTCCTTTGGAGAGGCTTACCCTCacagccatcatcaaagagatcgttag

For pRubi-PTEN-FBAR-T2A-Cre, srGAP3 in pCMV6 was ordered from Origene and Gibson assembly was used to fuse the FBAR domain in frame with PTEN using NEBuilder HiFi DNA Assemby Master Mix (NEB2621) and Q5 Hi-FI 2X Master Mix (NEBM0492) with the following:

pRubiPten-FBAR fwd gagacctaggGAGGGCAGAGGAAGTCTT pRubiPten-FBAR rev gagatgacatGACTTTTGTAATTTGTGTATGCTG pCMV6-entry SRGAP3_fwd tacaaaagtcATGTCATCTCAAACTAAGTTC pCMV6-entry SRGAP3_rev ctctgccctcCCTAGGTCTCCTCATTTTC PTEN-HOMER was a generous gift from Thomas Sudhof^71^. Gibson assembly was used to clone into pRubi-Pten-T2A-Cre with the following: PtenHOMER fwd agaatgcagcGAGGGCAGAGGAAGTCTTC Pten-HOMER rev ctcctccgccGACTTTTGTAATTTGTGTATGCTG Homer1c_fwd tacaaaagtcGGCGGAGGAGGCGGAGGC Homer1c_rev ctctgccctcGCTGCATTCTAGTAGCTTGGCCAAATTATCCCG

The Halo9 tag was subsequently cloned from pRubi-PTEN-HALO-T2A-Cre onto NLS-PTEN, NES-PTEN, PTEN-FBAR and PTEN-HOMER with the following:

Pten-FBAR_fwd gtcaaccctgGAGGGCAGAGGAAGTCTTC Pten-FBAR_rev cgctaccgccCCTAGGTCTCCTCATTTTCTGTG Halo9_fwd gagacctaggGGCGGTAGCGGGGATCCAC Halo9_rev ctctgccctcCAGGGTTGACAGCCATCTGGC Pten-Homer_fwd gtcaaccctgGAGGGCAGAGGAAGTCTTC Pten-Homer_rev cgctaccgccGCTGCATTCTAGTAGCTTGG Halo9_fwd agaatgcagcGGCGGTAGCGGGGATCCAC Halo9_rev ctctgccctcCAGGGTTGACAGCCATCTGGC 2xNLS-Pten_fwd gtcaaccctgTCTAGAGAGGGCAGAGGAAG 2xNLS-Pten_rev cgctaccgccGACTTTTGTAATTTGTGTATGCTGATCTTC Halo9_fwd tacaaaagtcGGCGGTAGCGGGGATCCAC Halo9_rev cctctctagaCAGGGTTGACAGCCATCTGGC NES-Pten_fwd gtcaaccctgTCTAGAGAGGGCAGAGGAAG NES-Pten_rev cgctaccgccGACTTTTGTAATTTGTGTATGCTGATC Halo9_fwd tacaaaagtcGGCGGTAGCGGGGATCCA Halo9_rev cctctctagaCAGGGTTGACAGCCATCTGG

### STEREOTAXIC INJECTION

Stereotaxic injection was performed as in^1,37,72^. Briefly, postnatal day (P) 7 mice were anesthetized with isoflurane and injected with 2 µL of virus per hemisphere at a rate of 0.3 µL/min at *y*+1.55mm and *x*±1.30mm relative to lambda. The *z*-coordinates were −2.3, −2.2, −2.1, −2.0 mm, with 25% of the total volume of virus injected at each depth. Upon completion of injection, two minutes were timed before slowly bringing the needle out of the skull.

### TRANSCARDIAL PERFUSION

At 21 days post-injection (DPI), mice were anesthetized with 2% 2,2,2-tribromoethanol (Sigma Aldrich) injected intraperitoneally. The anesthetized mice were transcardially perfused with cold filtered phosphate-buffered saline (PBS) with 4% sucrose for 5 minutes, followed by 4% paraformaldehyde (PFA, Sigma Aldrich) in PBS with 4% sucrose for 10 minutes. The collected brains were stored in 4% PFA in PBS with 4% sucrose for 24 hours following perfusion for post-fixation.

### FIXED-TISSUE IMMUNOHISTOCHEMISTRY

Brains were sliced coronally with a Leica VT1000S vibratome, and 50 µm, 100 µm, and 150 µm sections were collected depending on need. 50 µm sections were used to measure for localization, cross-sectional soma area, spine density, length, and head area, while 150 µm sections were used for dendritic arbor analysis. Immunohistochemistry was performed as described in^69,73^. Importantly, for anti-Pten IHC free-floating sections are permeabilized in PBS-T with 0.4% Triton X-100 for 30 minutes and Triton X-100 is not used in subsequent steps. Primary antibodies include chicken anti-GFP (1:3000; Abcam #13970), mouse anti-RFP (1:2000; Rockland #200-301-379), and rabbit anti-PTEN (1:80; Cell Signaling #9559). Secondary antibodies include Alexa488 goat anti-chicken (GFP; 1:200; Jackson Immunoresearch #103-545-155), Cy3 donkey anti-mouse (RFP; 1:200; Jackson Immunoresearch #715-165-150), and Alexa647 goat anti-rabbit (PTEN; 1:200; Jackson Immunoresearch #111-605-144).

### CONFOCAL IMAGING

Images were taken on a Zeiss LSM510 or LSM800 confocal microscope. Z-stacks for localization were imaged with a 40X oil immersion lens with 0.7x zoom, at 1024 x 1024 resolution, and a 2 µm z-step. Z-stacks for dendritic arborization were imaged with a 40X oil immersion lens with 0.7x zoom, at 1024 x 1024 resolution, and a 2 µm z-step. Z-stacks for soma size analysis were imaged with a 40x oil immersion/0.75 μm plan-apochromat lens with 0.7x zoom, at 512 x 512 resolution, and a 2-μm *z*-step. Z-stacks for spine density, length, and head area were imaged with a 63X oil immersion lens with 3x zoom, at 1024 x 1024 resolution, and a 0.5 µm z-step.

### CONFOCAL IMAGE ANALYSIS

Localization of PTEN was determined using fluorescence intensity ratios. Regions of interest (ROIs) were drawn with the ImageJ/Fiji (NIH) polygon tool. ROIs for soma data included the nucleus, a sliver of cytoplasm surrounding the nucleus, and background in the molecular layer. ROIs for the spine data included spine heads, dendrite, and background in the molecular layer. Fluorescence intensity was measured in ImageJ/Fiji (NIH), and ratios were calculated in Excel before transferring to GraphPad Prism.

Soma size was analyzed by an investigator blind to the experiment and experimental groupings, in the same manner as described in^73^, by circling somas at their maximum circumference in ImageJ/Fiji (NIH). Total dendritic arborization was analyzed by first collecting 150 µm sections from the hippocampus. *z*-Stacked images were taken on Zeiss LSM510 or LSM800 laser-scanning confocal at 40x oil immersion/0.75 plan-apochromat lens with 1x zoom, at 1024 x 1024, 4x averaging, and a 2-μm *z*-step. Neurons with cell bodies in the center 50 µm of the 150 µm image stacks were measured to ensure no significant arborization outside of the 150 µm thick section. Neurons were reconstructed, by a blinded investigator using user-guided, semi-automated Neurolucida 360 (MBF Biosciences), and analyzed with Neurolucida Explorer (MBF Biosciences)^37^.

Spine density, length, and head area of the first dataset were calculated using ImageJ/FIJI and Neuron Studio (Computational Neurobiology and Imaging Center at the Mount Sinai School of Medicine). Images were deconvolved into Richardson-Lucy tiffs using FIJI ^37^ and loaded into NeuronStudio which automatically quantifies spine density, length, and head area^74^.

We performed a second, independent spine head area analysis using the 150µm thick sections obtained for Neurolucida reconstructions. Due to the thickness of the sections 2photon microscopy was used to obtain high magnification images (Bruker 2P Plus) with a 20x 1.0NA water lens at 18x zoom using 1024×1024 images at 0.0434µm/pixel. Because the diffraction limit for emission in this setup is approximately 291nm, we estimated the spine head area using ImageJ/FIJI in 2 ways. The first was performed manually building an ROI around the head at its maximal circumference using the polygon tool, creating an intensity threshold, and quantifying the area of the spine based on thresholded pixels. We also analyzed randomized images in which a blinded investigator used the polygon tool to outline the perceived margins of the spine head, generating an ROI from which cross-sectional area was measured in FIJI.

### ACUTE SLICE GENERATION

Acute live slice generation was previously described in^37,75^. Briefly, mice were anesthetized with 2,2,2-tribromoethanol and perfused with an ice cold cutting solution containing the following (in mM): 110 choline-Cl, 10 D-glucose, 7 MgCl_2,_ 2.5 KCl, 1.25 NaH_2P_O_4-_2H_2O_, 0.5 CaCl_2_, 1.3 NA-ascorbate, and 25 NaHCO_3_, bubbled with 95% O_2-_5%CO_2._ 310 µm-thick sections were generated through the hippocampus on a VT1200s vibratome (Leica), For 30 minutes post-slicing, slices were stored in artificial cerebrospinal fluid (aCSF) at 34°C, then moved to room temperature. The contents of aCSF are as follows (in mM): 125 NaCl, 25 NaHCO_3,_ 2.5 KCl, 1.25 NaH_2P_O_4_, 2.0 CaCl_2_, 1.0 MgCl_2_, and 25 D-glucose, bubbled with 95% O_2-_5%CO_2._ Slices were incubated with 100nM JF-669-Halo in ACSF for 1h followed by a rinse and 30 minute wash prior to imaging using continuous ACSF flow.

### LIVE-TISSUE MULTI-PHOTON IMAGING

Multi-photon microscopy (Bruker Ultima 2p Plus) was utilized to capture a *z*-series (0.5 µm step) through the entire dendritic arbor of retrovirus-infected granule neurons every 1 minute for 30 mins from acute slices collected between 7- and 12-days post-injection (DPI). Neurons were imaged with 20x 1.0NA lens, with 4x averaging, 4x zoom, at either 512×512 pixels or 1024×1024 pixels, and a 1.2 µs dwell time. Slices were continually bathed with aCSF bubbled with 95% O_2_, 5% CO_2_ at 37°C. Slices were incubated with 100nM JF669 for 1h followed by at least 30min washout. Fluorescence was stimulated using 1040nM for tdTomato and 1240nM for JF-669 using a Chameleon Discovery NX tunable laser (Coherent).

### LIVE-TISSUE MULTI-PHOTON IMAGE ANALYSIS

Images were analyzed in ImageJ/Fiji. Images were first converted from a stack to a hyperstack, and channels were separated into cyan (Pten-Halo) and red (tdTomato cytoplasm). The Fiji smooth algorithm was used reduce single pixel noise. The n-Tracer1.1.1 alignmaster plug-in^76^ was utilized to correct for photobleaching over time. Finally, channels were merged, and the SIFT MultiChannel plugin was utilized to control for *x*- and *y*-drift.

### PULSE-CHASE

Wild type mice (Jackson Laboraotries; C57BL/6J) mice were injected with 2 µL of virus per hemisphere at a rate of 0.3 µL/min at *y* + 1.55 mm and *x* ± 1.30 mm relative to λ. The *z*-coordinates were −2.3, −2.2, −2.1, −2.0 mm, with 25% of the total volume of virus injected at each depth. Upon completion of injection, we waited 2 minutes before slowly bringing the needle out of the skull to ensure full saturation of virus. The virus used was AAV-DJ-hSyn-PTEN-Halo9-T2A-Cre. Halotag dye JFX673 and JF552 were generously gifted from the Lavis lab at Janelia Research Campus. Both dyes were prepared the day before usage by adding 40 µL DMSO (Invitrogen; D12345), 40 µL Pluronic F-127 (Invitrogen; P3000MP), and 120 µL of 0.9% Bacteriostatic NaCl injection saline containing benzyl alcohol (Hospira; NDC0409-1966-02), then wrapping in foil and placing onto a shaker at 200 RPM at 25C overnight. At 21DPI, mice were given a 50 µL “pulse” dose retro-orbitally of JFX673 HaloTag dye (; CN-1-170) to label HaloTag domain of the PTEN. After a variable amount of time a 50 µL “chase” dose was given retro-orbitally of JF552 HaloTag dye (Lavis Lab; SEP-4-035) to tag any newly synthesized PTEN protein. The pulse-chase intervals were: 1.5h, 8h, 24h, 48h, 72h, 96h. Mice were then anesthetized using 2% Avertin and perfused with ice-cold 4% PBS-sucrose then fixed using ice-cold 4% PFA. Brains were then extracted and put into 4% PFA for 24 hours before being switched into 4% PBS-sucrose. Tissue was then sectioned using a vibratome at 100 um slices and collected for mounting. Slices were mounted and coverslipped using hard mounting media without DAPI (Vectashield; H-1400-10) and allowed to cure overnight before imaging. Images for quantification were taken using multi-photon microscopy (Bruker Ultima 2p Plus) with a 20x 1.0NA lens, with 4x averaging, 4x zoom, at 1024×1024 pixels and a 0.6 µs dwell time. Fluorescence was stimulated at 850 nm at 160 power, using a Chameleon Discovery NX tunable laser (Coherent). Each slice was imaged for the hilus of the DG, the granule and molecular layers of the DG and the mossy fiber axonal projections from the DG to the CA3 of the hippocampus proper.

To quantify the lifetime of PTEN from the images, the fluorescence intensity of the pulse and chase dyes were taken using FIJI, then the intensities were used to calculate the mean lifetime with the formula below^77^. In this formula, Δt is the interval between the pulse and chase injections, I is the mean fluorescence intensity of each dye, and ln is the natural logarithm. To assess whether PTEN lifetime varied across different areas of the neuron itself, images from the granule and molecular layer of the DG were split into ROIs consisting of the somal bodies of the neuron, dendrites of the neuron and then the entire region combined. Images were thresholded to minimize background noise.

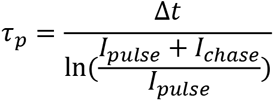

Statistical significance between regions was assessed using a one-way ANOVA and post-hoc analysis was done using a Tukey’s multiple comparison test.

### EXPERIMENTAL DESIGN AND STATISTICAL ANALYSIS

Quantitative values reported in the manuscript text are mean±SEM (n=animals unless otherwise indicated). Wildtype and *PTEN* KO data from Figure 4P, Figure 5H-M, and Figure 6H-J is a combination of new data and data previously published ^78^. No differences were found between previously generated and current control data. All graphs were generated using GraphPad Prism with the specific test indicated in figure legends. Unless otherwise indicated, statistical comparisons were performed using a mixed effects model accounting both for the number of animals and neuronal measurements clustered within animals using Stata ^79^. For examining the frequency distribution of *Pten* between the cytosol and nucleus or dendrite and spine the Kolmogorov-Smirnov test was used per cell (cytoplasm/nucleus) or per spine (spine/dendrite). Fit lines for the distribution were generated using the Asymmetric Sigmoidal, 5 parameter logistic, X is log(concentration). For Sholl analysis statistical comparisons were performed using the mixed effects model with Tukey’s multiple comparisons.

## Supplemental Video Legend

**Supplemental Video 1.**

A P7 *Pten*^flx/flx^ x Ai14 mice was infected with PTEN-Halo9-T2A-Cre and acute slices are generated for live 2-photon imaging at 9 DPI. PTEN-Halo9 was visualized using JF669 excited at 1240nm and tdTomato at 1040nm. A 29-image z-stack with a 0.5µm step size was captured every 2 minutes for 1 hour. The movie first shows the maximum projection of PTEN-Halo9 (cyan) and tdTomato (red) with balanced brightness and contrast followed by the entire movie with the balance and contrast optimized for PTEN-Halo9 (cyan), then tdTomato (red).

**Supplemental Video 2.**

Another example 2-photon movie for P7 *Pten*^flx/flx^ x Ai14 mice are infected with PTEN-Halo9-T2A-Cre and acute slices are generated for live 2-photon imaging at 8 DPI. A 25-image z-stack with a 0.5µm step size was captured every 2 minutes for 1 hour. Frames 1-30 show the maximum projection of PTEN-Halo9 (cyan) and tdTomato (red) with balanced brightness and contrast followed by all frames with the balance and contrast optimized for PTEN-Halo9 (cyan), then tdTomato (red).

## Notes

https://neuromorpho.org/index.jsp

### Competing Interest Statement

The authors have declared no competing interest.

### Summary of Updates

The PTEN lifetime values reported from the DELTA pulse-chase experiment have been corrected. In the original analysis the mean lifetime was calculated with a base-10 logarithm. The DELTA method (Mohar et al., 2025) defines the mean lifetime with the natural logarithm, tau = t / ln(1/fraction pulse). As a result, every lifetime value in the original version was too large by a constant factor of ln(10), or 2.30. The corrected average lifetime of PTEN-Halo9 in dentate granule neurons is 2.50 plus or minus 0.14 days (previously reported as 5.76 plus or minus 0.32 days). The corrected value is reported in the Results, the Discussion, and the legend of Figure 2. Figure 2H has been replotted with the corrected values. The Methods now state the formula explicitly and define each term, including the natural logarithm. The relative comparisons in the paper are unchanged because the correction is a constant factor. The comparison to other proteins measured with DELTA has been updated. PTEN lifetime is roughly half that of GluA2 and much shorter than PSD-95 and MeCP2. This strengthens our interpretation that PTEN is a relatively unstable and highly regulated protein. No other data, analyses, or conclusions were changed.

